# hilldiv: an R package for the integral analysis of diversity based on Hill numbers

**DOI:** 10.1101/545665

**Authors:** Antton Alberdi, M Thomas P Gilbert

**Author notes:** **Correspondence**: Antton Alberdi Øster Farimagsgade 5, KH7, DK-1353 Copenhagen, Denmark.

## Abstract

Hill numbers provide a powerful framework for measuring, comparing and partitioning the diversity of biological systems as characterised using high throughput DNA sequencing approaches. In order to facilitate the implementation of Hill numbers into such analyses, whether focusing on diet reconstruction, microbial community profiling or more general ecosystem characterisation analyses, we present a new R package. ‘Hilldiv’ provides a set of functions to assist analysis of diversity based on Hill numbers, using count tables (e.g. OTU, ASV) and associated phylogenetic trees as inputs. Multiple functionalities of the library are introduced, including diversity measurement, diversity profile plotting, diversity comparison between samples and groups, multi-level diversity partitioning and (dis)similarity measurement. All of these are grounded in abundance-based and incidence-based Hill numbers, and can accommodate phylogenetic or functional correlation among OTUs or ASVs. The package can be installed from CRAN or Github, and tutorials and example scripts can be found in the package’s page (https://github.com/anttonalberdi/hilldiv).

## Introduction

Tools for analysing diversity lie at the core of molecular ecology. For example, whether profiling dietary content, microorganism communities or any other bulk samples using genetic tools, researchers routinely need to compare diversity between samples, partition diversity between different hierarchical levels, and/or compute (dis)similarity measures between samples. Although a wide repertoire of metrics have been developed to perform such operations, there is an increased awareness of the need to use general statistical frameworks to generate results that are more consistent and more easily interpretable (Jost, 2006; Chao, Chiu, & Jost, 2010).

One such general statistical framework that can enable diversity analysis is that developed around the so-called ‘Hill numbers’ (Hill, 1973; Jost, 2006). This framework provides a robust toolset with which to perform the most common operations researchers routinely use when analysing the diversity of biological systems, and includes among others, diversity measurement and estimation, diversity partitioning, diversity decomposition and (dis)similarity computation (Alberdi & Gilbert, 2019). R packages containing functions to perform basic diversity analyses based on Hill numbers already exist, including *entropart* (Marcon & Hérault, 2015), *vegan* (Oksanen et al., 2013), *vegetarian* (Charney & Record, 2012), *hillR* and *simba* (Jurasinski, 2007). While these tools have enabled incorporating Hill numbers into the statistical toolbox of molecular ecologists, the usage of Hill numbers is usually limited to diversity measurement and estimation (Hsieh, Ma, & Chao, 2016). One underlying reason might simply be the lack of easy-to-use tools that compile functions for the integral analysis and visualisation of a wide range of diversity analyses based on Hill numbers.

To meet this need, we present ‘hilldiv’, an R package that encompasses different functions with which to perform a number of diversity analyses based on Hill numbers. Although it’s potential uses are wider, ‘hilldiv’ is primarily designed for diversity analyses of molecularly (e.g. metabarcoding) characterised datasets, using count tables of OTUs and ASVs (hereafter just OTUs for the sake of readability) and related phylogenetic trees. The package includes functions for diversity measurement, diversity profile generation and plotting, diversity comparison between samples and groups, diversity partitioning at multiple hierarchical levels and (dis)similarity measurement among others (Fig. 1).

**Figure 1.**
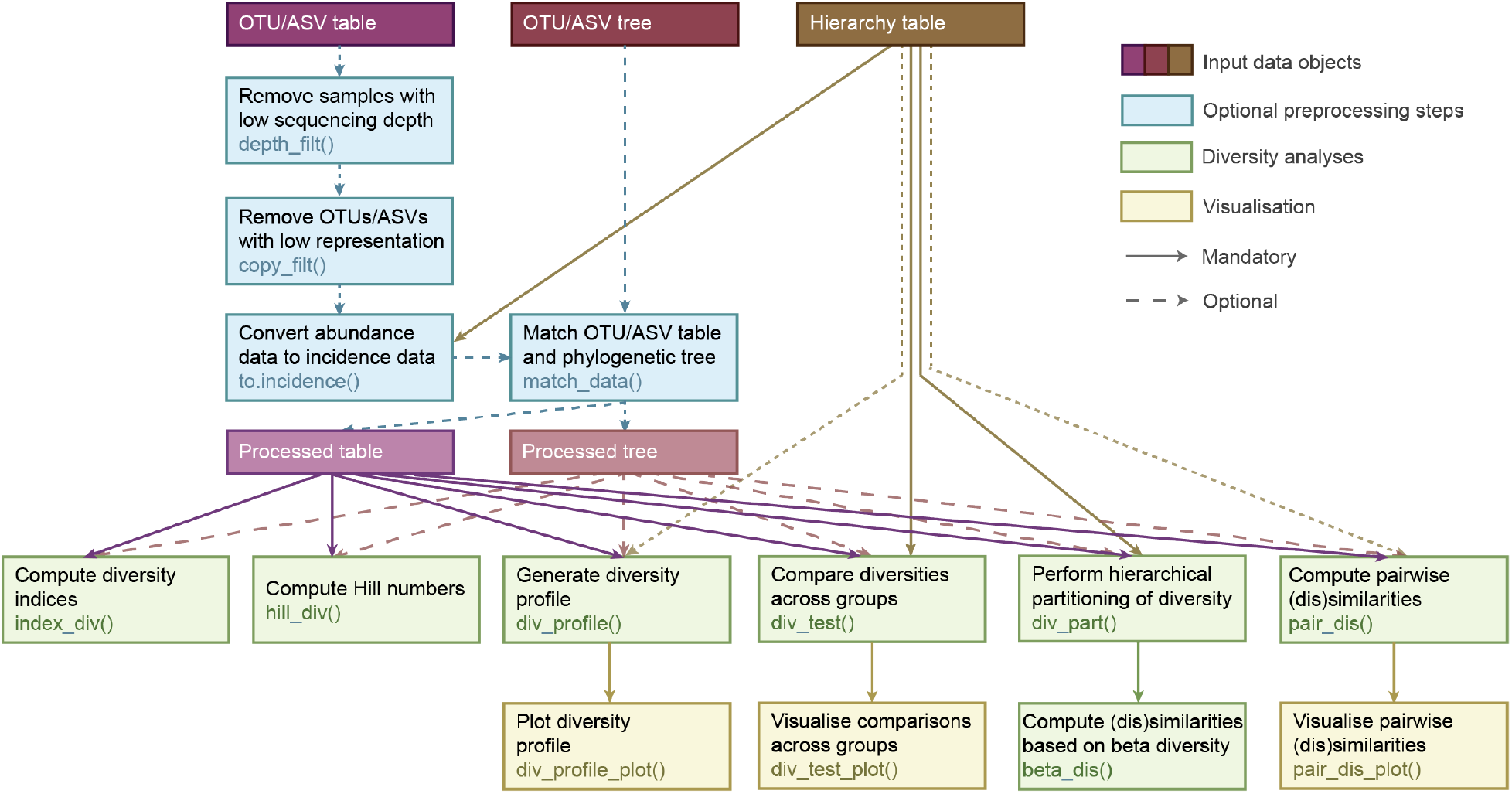
**Schematic representation** of the nature and relations across the main functions included in ‘hilldiv’.

## Statistical background

The statistical framework developed around Hill numbers encompasses (through monotonic transformations) many of the most broadly employed diversity (e.g. richness, Shannon index, Simpson index), phylogenetic diversity (e.g. Faith’s PD, Allen’s H, Rao’s quadratic entropy) and dissimilarity (e.g. Sørensen index, Unifrac distances) metrics (Chiu, Jost, & Chao, 2014). This enables the most common analyses of diversity to be performed while grounded in a single statistical framework, and provides a number of benefits in comparison to the use of each of the other metrics separately:

i. Hill numbers meet the so-called doubling property, which means that when doubling the number of OTUs in a system while maintaining the rest of the parameters (e.g. evenness, phylogenetic relations between OTUs), then the diversity measured is also doubled (Chao et al., 2010). This is a property that richness owns, but Shannon and Simpson indices, for instance, do not (Jost, 2006).
ii. The interpretation of both the measure and measurement unit is consistent for each type of data. The basic Hill numbers expression yields a diversity measure in “effective number of OTUs”, which is interpreted as the number of equally abundant OTUs that would be needed to give the same value of diversity. This contrasts, for instance, with Shannon and Simpson indices (Shannon, 1948), which yield uncertainty and probability values, respectively (Jost, 2006).
iii. The sensitivity towards abundant and rare OTUs can be modulated with a single parameter, namely the order of diversity (q). When a diversity of order one (q=1) is used, the OTU relative abundances are weighed as their original values. However, when a diversity of order q<1 is used, rare OTUs are overweighed. When taken to its extreme (diversity of order set to zero, q=0), relative abundances are not considered at all, and the data simply reflects presence/absence. In contrast, when orders of diversity of q>1 are used, abundant OTUs are overweighed. Three q values are particularly relevant, for their close relationship to popular diversity indices. The Hill number of q=0 yields a richness value, the Hill number of q=1, also known as Shannon diversity, is the exponential of the Shannon index, and the Hill number of q=2, also known as the Simpson diversity, is the multiplicative inverse of the Simpson index (Jost, 2006).
iv. Hill numbers can also be computed while considering the phylogenetic relationships among OTUs. When doing so, the phylogenetic Hill number of q=0 equals Faith’s PD divided by tree depth, the phylogenetic Hill number of q=1 is the exponential of Allen’s H divided by tree depth, and the phylogenetic Hill number of q=2 is the multiplicative inverse of Rao’s Q divided by tree depth.
v. Hill numbers can be computed from both abundance and incidence data. In abundance-based approaches the DNA sequence is the unit upon which diversity is computed, while in incidence-based approaches the sample is the unit upon which diversity is measured (Chao et al., 2014). Therefore, abundance-based Hill numbers measure the effective number of equally abundant OTUs in the system, while incidence-based Hill numbers measure the effective number of equally frequent (across samples) OTUs in the system (Chao et al., 2014).
vi. The Hill numbers framework enables the diversity of a system to be partitioned into multiple hierarchical levels following the multiplicative definition (Jost, 2007; Chao, Chiu, & Hsieh, 2012).
vii. It is possible to compute multiple (dis)similarity measurements derived from beta diversities, both for Hill numbers and phylogenetic Hill numbers.

## Applications

Version 1.5.0 of the ‘hilldiv’ package contains 25 functions that enable different types of diversity measurements to be performed, as well as compared and visualised. The package is structured in a way that enables non-experienced R users to perform statistical tests and generate plots in a straightforward way, while allowing experienced R users to obtain raw results as vectors, matrices or lists to perform advanced operations with other packages (Fig. 1). For operating some of the scripts, ‘hilldiv’ calls functions from other R packages, including *ape* (Paradis, Claude, & Strimmer, 2004), ggpubr (Kassambara, 2017), *ggplot2* (Wickham, 2010), qgraph (Epskamp et al., 2012) and *vegan* (Oksanen et al., 2013).

In the following, we introduce the main functions for performing the most relevant tasks. Simple examples of the most relevant functions are also provided, while more complex examples can be found in the package’s documentation page: https://github.com/anttonalberdi/hilldiv/wiki.

### Data

The package contains three example datasets that enable all of its functions to be reproduced. The OTU table is a *dataframe* that contains Illumina amplicon read abundance data (40 samples and 363 OTUs) derived from faecal samples generated by (Alberdi, Aizpurua, Gilbert, & Bohmann, 2018). The OTU phylogenetic tree is a *phylo* object that contains the phylogenetic relations across OTUs. The hierarchy table is a *dataframe* that contains the relation between sample names and their respective parent group, in this case taxonomic species.

Example 1: load example data sets.

~~~
data(bat.diet.otutable)
data(bat.diet.tree)
data(bat.diet.hierarchy)
~~~

### (Phylo)diversity computation

The diversity of individual samples can be calculated for vectors (one sample) or matrices (OTU table containing multiple samples) using the function *hill_div()*. The function requires as input the abundance data (as a vector or matrix), and specification of the order of diversity (q-value) at which diversity will be computed. As default, *hill_div()* yields the diversity measure in ‘effective number of OTUs’. If an ultrametric tree object that establishes the (e.g. phylogenetic) relations between OTUs is also provided, the function considers the correlation between OTUs, and yields the diversity measure in ‘effective number of lineages’. The package also contains the function *index_div()*, that enables the six most common diversity indices associated to Hill numbers to be computed: richness, Shannon index, Simpson index, Faith’s PD, Allen’ H and Rao’s Q.

Example 2: phylogenetic Hill numbers of order of diversity 1 of all samples in the OTU table.

~~~
hill_div(bat.diet.otutable, qvalue=1, tree=bat.diet.tree)
~~~

Example 3: Allen’s H of all samples in the OTU table.

~~~
index_div(bat.diet.otutable, tree=bat.diet.tree, index="allen")
~~~

### (Phylo)diversity profile

An effective way for representing the different components of the diversity of a system is to plot its diversity profile, as it provides information about the richness and evenness of a sample at a glance (Fig. 2a) (Chiu et al., 2014). This can be done by generating the data using *div_profile()* and plotting it using *div_profile_plot()*. These two functions enable generating and plotting diversity profiles for a single sample (Fig. 2b), a set of samples (Fig. 2c) or multiple sets of samples aggregated by groups as specified in the hierarchy table (Fig. 2d). If a hierarchy table is provided, it is possible to generate and plot either the alpha or gamma diversity profiles of the groups as specified by the argument ‘level’.

**Figure 2.**
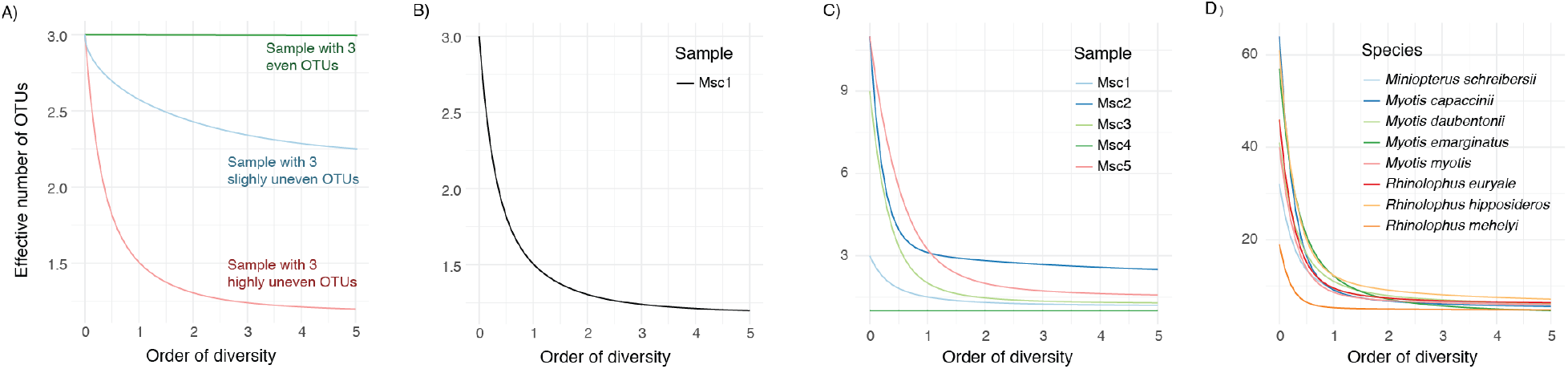
Diversity profiles. A) Three diversity profiles of hypothetical data sets with 3 OTUs with different abundance distributions, and B-D) diversity profiles generated with the function *div_profile()* for B) one sample, C) multiple individual samples, and D) multiple samples aggregated in groups.

Example 4: generate and plot the gamma diversity profiles of the groups specified in the hierarchy table.

~~~
profiledata <-div_profile(bat.diet.otutable,hierarchy=bat.diet.hierarchy)
div_profile_plot(profiledata)
~~~

### (Phylo)diversity comparison tests

Researchers often need to compare diversity among different groups of samples. This can be easily achieved using the function *div_test()*, which performs statistical hypothesis testing between the Hill numbers of the groups specified in the hierarchy table. The function assesses normality and variance homogeneity of the data, and depending on that performs either Student’s t-test or Wilcoxon Rank Sum Test if there are 2 groups, and either ANOVA or Kruskal-Wallis test if there are multiple groups. The function also enables pairwise post-hoc analyses to be performed. The related function *div_test_plot()* uses the output object of the *div_test()* function to visually summarise the diversity values (Fig. 3A), including the option to show p-values of pairwise comparisons (Fig. 3B).

**Figure 3.**
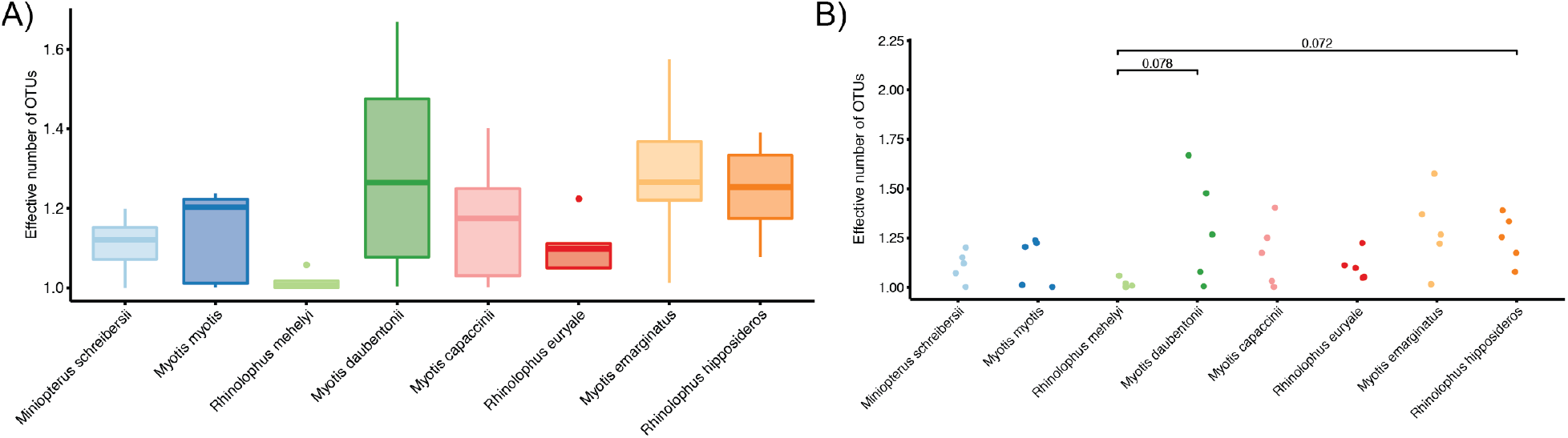
**Diversity comparison plots** produced using the *div_test_plot()* function. A) Box plot generated using the chart argument “box”, B) Jitter plot generated using the chart=“jitter”, posthoc=TRUE and threshold=0.1.

**Figure 4.**
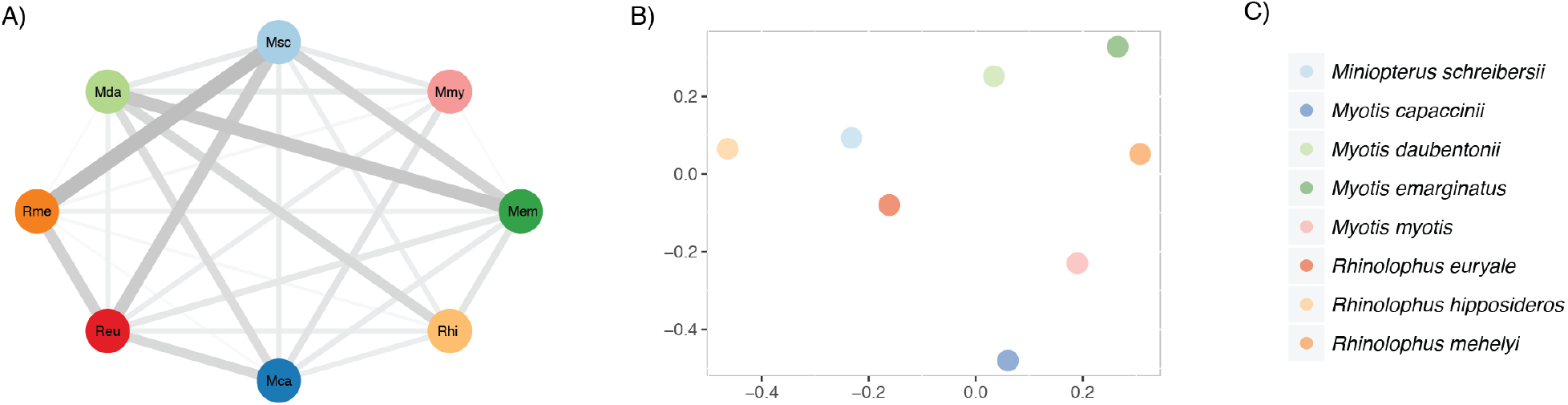
Dissimilarity visualisation plots. A) A qgraph network diagram, B) a NMDS chart and C) legend of the groups.

Example 5: contrast the sample-level phylogenetic Hill numbers across different species including pairwise post-hoc analyses and plot the results as a jitter plot showing pairwise p-value below p=0.1.

~~~
divtestresult <-div_test(bat.diet.otutable,qvalue=1,hierarchy=bat.diet.hierarchy,tree=b at.diet.tree,posthoc=TRUE)
div_test_plot(divtestresult, chart="jitter", posthoc=TRUE, threshold=0.1)
~~~

### (Phylo)diversity partitioning

The function *div_part()* enables diversity partitioning into different hierarchical levels following the multiplicative definition to be performed. If no hierarchy table is specified, the functions partitions the diversity into sample-level (L1 = alpha) and overall system (L2 = gamma) diversities, and also yields the corresponding L1_2 (beta) value. If a hierarchy table is provided, the function yields diversities of all specified levels (L1, L2, L3 … Ln) and their respective beta values (L1_2, L2_3 … L(n-1)_n). The structure specified in a hierarchy table with more than three levels must be nested, i.e. two samples that belong to a common parent group cannot have different grandparent groups.

Example 6: partitioning of phylogenetic diversity of order of diversity 0 into two levels.

~~~
div_part(bat.diet.otutable,qvalue=0,tree=bat.diet.tree)
~~~

Example 7: partitioning of diversity of order of diversity 1 into more than two levels.

~~~
div_part(bat.diet.otutable,qvalue=1,hierarchy=bat.diet.hierarchy)
~~~

### (Dis)similarity measurement

The function *beta_dis()* performs dissimilarity or similarity measurement based on Hill numbers beta diversity, sample size and order of diversity. The function can be run by inputting those values manually, or by using the object outputted by the *div_part()* function, which contains all the mentioned information. As specified by the argument “metric”: the function can compute the following similarity measures: the Sørensen-type overlap (C_qN_), the Jaccard-type overlap (U_qN_), the Sørensen-type turnover-complement (V_qN_), and the Jaccard-type turnover-complement (S_qN_). The argument ‘type’ enables either similarities or dissimilarities (one-complements of the similarity values) to be outputted.

Example 8: computing Sørensen-type overlap from manually inputted data.

~~~
beta_dis(beta=4, qvalue=1, N=8, metric="C", type="similarity")
~~~

Example 9: computing Jaccard-type turnover at the sample (L1_2) and species (L2_3) level from multi-level diversity partitioning of order of diversity 1.

~~~
divpartres <-div_part(bat.diet.otutable,qvalue=1,hierarchy=bat.diet.hierarchy) beta_dis(divpartres, metric="S", type="dissimilarity")
~~~

### Pairwise (dis)similarity

The function *pair_dis()* performs pairwise diversity partitioning and yields matrices containing pairwise beta diversity and corresponding (dis)similarity measures. If a hierarchy table is provided, pairwise calculations can be carried out at some or all the specified hierarchical levels. The results are outputted as a list of matrices. The related function *pair_dis_plot()* uses any of the dissimilarity matrices yielded by *pair_dis()* (e.g. 1-U_qN_) to visualize it either as a qgraph plot or a NMDS chart.

Example 10: computation of the Jaccard-type overlap-complement among species (L2: second hierarchical level specified in the hierarchy table)

~~~
pairdisres <-pair_dis(bat.diet.otutable, qvalue=0, hierarchy=bat.diet.hierarchy,level=2,metric="U")
~~~

Example 11: visualisation of the Jaccard-type overlap-complement as a qgraph or a NMDS chart

~~~
pair_dis_plot(pairdisres$L2_UqN, hierarchy=bat.diet.hierarchy, type="qgraph",level=2)
pair_dis_plot(pairdisres$L2_UqN, hierarchy=bat.diet.hierarchy, type="NMDS",level=2)
~~~

### Auxiliary functions

The package also includes a range of auxiliary functions. The function *to.incidence()* enables conversion of an abundance-based count table into an incidence-based table, so that the aforementioned functionalities can also be implemented based on incidence data. The function *match_data()* compares the count table with the phylogenetic tree and removes the OTUs missing in one of the two data sets. The functions *depth_filt()* and *copy_filt()* enable removing samples and OTUs, respectively, based on user-specified minimum depth or representation levels.

Example 12: convert count table containing abundances into an incidence table based on the hierarchy table.

~~~
bat.diet.otutable.inc <-to.incidence(bat.diet.otutable, bat.diet.hierarchy)
~~~

## Conclusions

The package ‘hilldiv’ enables many of the most common statistical operations that molecular ecologists routinely require to be performed in a straightforward way within the statistical framework developed around the Hill numbers. Although it is primarily devised for metabarcoding data containing multiple samples and OTUs, it could also be useful to perform diversity analyses on shotgun metagenomics data, or indeed any other non-molecular data that is comprised of multiple sampling units and types that enable measures of diversity to be quantified.

## Data accessibility

All the data used in this paper are included in the R package. The R package is currently available at CRAN and Github (https://github.com/anttonalberdi/hilldiv).

## Author contributions

A.A and M.T.P.G devised the project. A.A wrote the functions and drafted the manuscript. Both authors contributed to the writing of the manuscript.

## Acknowledgements

A.A. was supported by Lundbeckfonden (grant R250-2017-1351) and M.T.P.G. acknowledges ERC Consolidator Grant (681396-Extinction Genomics).

